# Epigenetic fidelity in complex biological systems and implications for ageing

**DOI:** 10.1101/2023.04.29.538716

**Authors:** Thomas Duffield, Laura Csuka, Arda Akalan, Gustavo Vega Magdaleno, Ludovic Senez, Daniel Palmer, João Pedro de Magalhães

**Author notes:** Correspondence to: Thomas Duffield or João Pedro de Magalhães.

## Abstract

The study of age is plagued by a lack of delineation between the causes and effects within the ageing phenotype. This has made it difficult to fully explain the biological ageing process from first principles with a single definition. Lacking a clear description of the underlying root cause of biological age confounds clarity in this critical field. In this paper, we demonstrate that the epigenetic system has a built-in, unavoidable fidelity limitation and consequently demonstrate that there is a distinct class of DNA methylation loci that increases in variance in a manner tightly correlated with chronological age. We demonstrate the existence of epigenetic ‘activation functions’ and that topological features beyond these activation functions represent deregulation. We show that the measurement of epigenetic fidelity is an accurate predictor of cross-species age and present a deep-learning model that predicts chronological age exclusively from knowledge of variance. We find that the classes of epigenetic loci in which variation correlates with chronological age control genes that regulate transcription and suggest that the inevitable consequence of this is a feedback cycle of system-wide deregulation causing a progressive collapse into the phenotype of age. This paper represents a novel theory of biological systemic ageing with arguments as to why, how and when epigenetic ageing is inevitable.

## Introduction

Despite increased research and the undeniable importance and impact of ageing in medicine and society (1), the exact nature of human ageing and its causative mechanisms remain largely controversial. Many theories have been put forward attempting to explain the ageing process (2), yet the underlying molecular drivers of the human ageing process continue to be a subject of great interest and intense debate (3).

Recent studies have put the limelight on the potential role of epigenetic modifications in ageing (4; 5; 6). These include the discovery of epigenetic clocks, highly accurate predictors of chronological age, based on a relatively small number of methylation sites (7; 8). Epigenetic clocks are associated with mortality, they can predict chronological age from various tissues, across the lifespan and in multiple species, although their mechanistic basis remains the subject of debate (5; 9). In addition, multiple changes in methylation and other epigenetic modifications have been reported with age, both in human and animal models (4; 6; 10).

It has been proposed that epigenetic changes are causative in ageing (5), and a recent study has suggested that DNA damage response-induced loss of epigenetic information drives ageing (11). More broadly, the information theory of ageing has suggested that loss of epigenetic information with age is a major driver of the ageing process (12; 11). It has also been suggested that pre-programmed shifts in epigenetic information states with age are a major determinant of ageing phenotypes (13). As such, understanding the basis of epigenetic clocks, and how epigenetic changes could impact ageing is a major and important open question. Moreover, despite efforts to understand the informatic character of ageing, there has been comparatively little research on what makes mammalian ageing inevitable.

In this work, we develop a conceptual model to explain the ageing process based on first principles. We demonstrate that the epigenetic system has unique inherent informatic properties that progressively acquire informatic corruption, meaning that with age epigenetic information fidelity cannot be maintained. Our model is further supported by empirical data from humans and other species, and we derive a predictor of age based solely on measures of epigenetic variation.

### 1 The fidelity limit theory of age

#### Repairing Damaged Information

Any state, including that of DNA methylation, can be thought of as a state of information(Fig 1(ii)), and therefore epigenetic damage (which we define as any epigenetic change that reduces the organism’s overall chance of survival, and thus is selected towards system ontology) represents information loss. When information is destroyed, there are only two possible mechanisms by which it can be recovered(Fig 1(iii)). Information in the original state can be recovered from an identical backup, via a system encoded to know which is which. Alternatively, information can be reconstituted through an algorithm: a series of rules that defined the original information state. All of these systems of data recovery must be applied consequently to damage through either observation or prediction of data loss.

**Figure 1:**
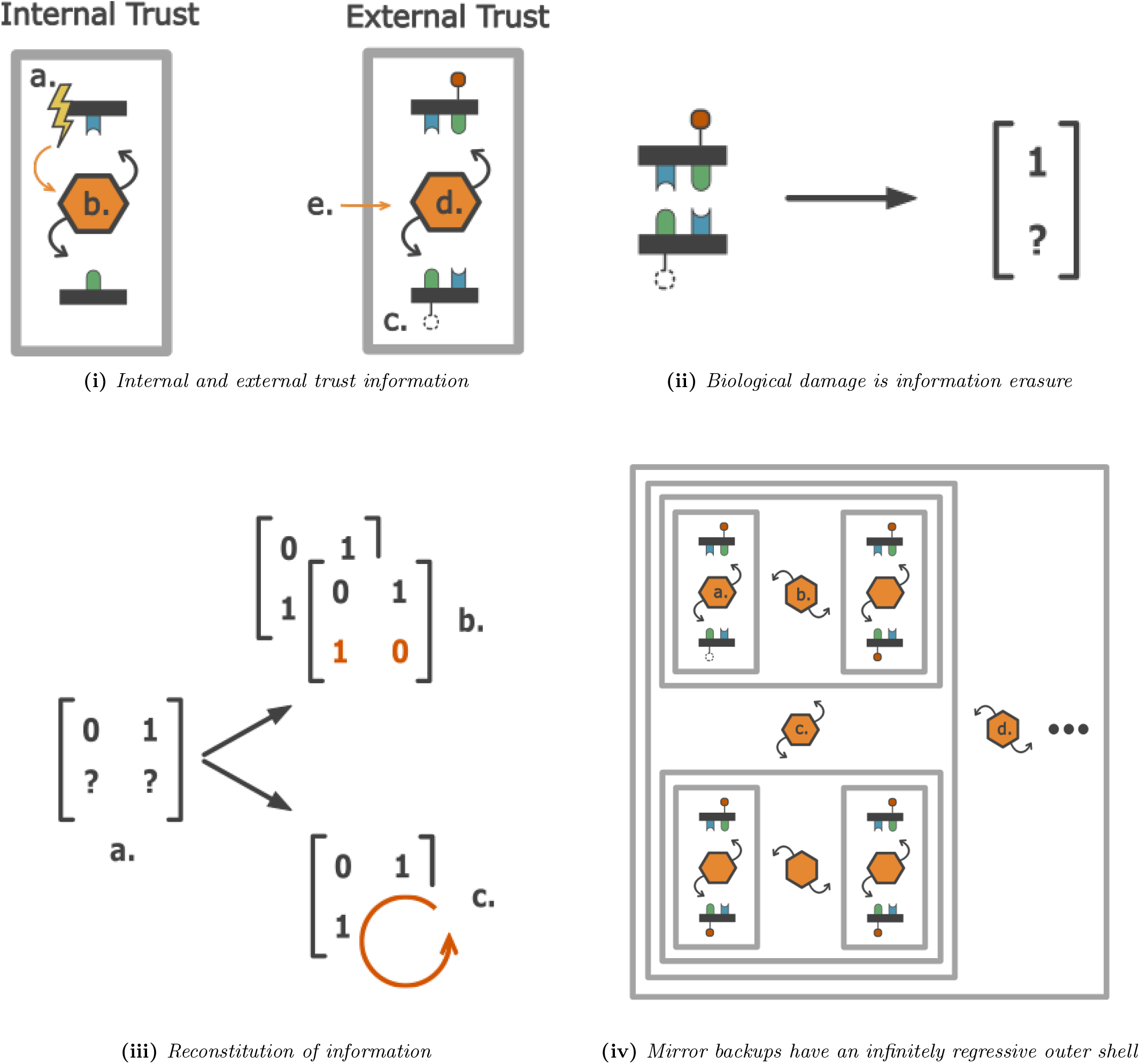
i. Natural selection infers that in single-strand breaks the side with the broken backbone (a.) is most likely to contain the incorrect base. A system for deducing which strand to use as a mirror backup (b.) will have access to this information. Methylation damage (c.) cannot provide this information, so (d.) would require information from an external scope (e.) to make the same comparison. ii. Incorrect modification of CpG methylation can be thought of as information erasure, removing part of the state information that allows for the recovery of the original state. iii. Epigenetic damage is information loss (a.) that requires repair either with a mirror backup from which to duplicate information (b.) or an algorithm with which to define it according to original principles(c.). iv. Epigenetic mirrored backups represent a vicious infinite regress of endlessly nesting scopes. Comparing two strands requires a system of trust recognition (a.) that, if subject to noise, would itself require a mirror backup. These two systems would themselves require an external system of trust (b.), which can itself make errors, requiring a backup and another system of trust (c.) and so on (d.)

### Mirror backup is impossible

To create a mirror backup for DNA methylation, an object with identical informatic properties would have to exist from which to duplicate the information, which would also have to behave in an identical manner to the first in response to noise and errors. Were this not the case, there would no longer be one-to-one parity between original and backup, and the process of comparison would itself become noisy. A system would then be necessary to duplicate changes between original and backup, maintaining parity, encoding trust, and indicating which of the two DNA methylation signals should be treated as a backup in the event of damage. In a biological context, such a system would itself inevitably be subject to error, allowing noise to enter the decision-making governing trust and therefore requiring another mechanism for mitigation and correction. In essence, just as DNA methylation is the single outer layer of control for DNA, DNA methylation itself would require the same system, which would be subject to the exact problems it was intended to avoid (Fig 1(iv)). This “nesting doll problem” is infinitely recursive: it is logically impossible in a noise-filled environment to design a signal without a component in which all damage has a mirror backup. We suggest a flawless epigenetic mirror is impossible as an example of vicious infinite regress (14), extremely similar to Bradley’s regress (15; 16). Although described here in terms of individual methylation loci, this process holds true for regions of methylation or even systems of comparison between chromosomes. Any such comparison requires a ‘comparer’, which becomes the point of entry for signal corruption, unless it itself has a backup and so on.

### Algorithmic fidelity is restricted

Lacking a mirror backup, any information lost in epigenetic damage must be reconstituted using some form of algorithm. Any algorithm that reconstitutes information must itself be encoded which, in the context of the cell, means genetically encoded in DNA. This has a consequent cost to the cell (for example, the more DNA used, the greater the chance of mutation), meaning that any increase in survival cost must be offset with additional functionality. Minimum algorithm size increases with the complexity of information it is to define: an increase in the latter must result in an increase in the former(Fig 2(i)). This means algorithm size is also related to the fidelity by which it reconstitutes lost information because low fidelity reconstitution represents a reduction in information from the original (Fig 2(ii)), essentially performing lossy compression (17). A perfect reconstitution requires the exclusive use of lossless compression and has consequently higher storage requirements. Natural selection will not select for lossless compression if the cost of the additional information outweighs the benefit to survival, meaning in all cases one should expect DNA compression to be lossy (except in the case of individual errors with infinite cost to survival, e.g. errors leading to cancer). With DNA methylation containing two legal character states, defining it with perfect fidelity would be equivalent to binary key definition in cryptology, becoming exponentially large as regions contain more CpG loci. With approximately 20 million CpG in the human genome, perfect fidelity is therefore impossible. Even working under the assumption that epigenetic regions represent the states to define, there are over 20000 CpG islands in the human genome and an uncountable number of cellular identities to define.

**Figure 2:**
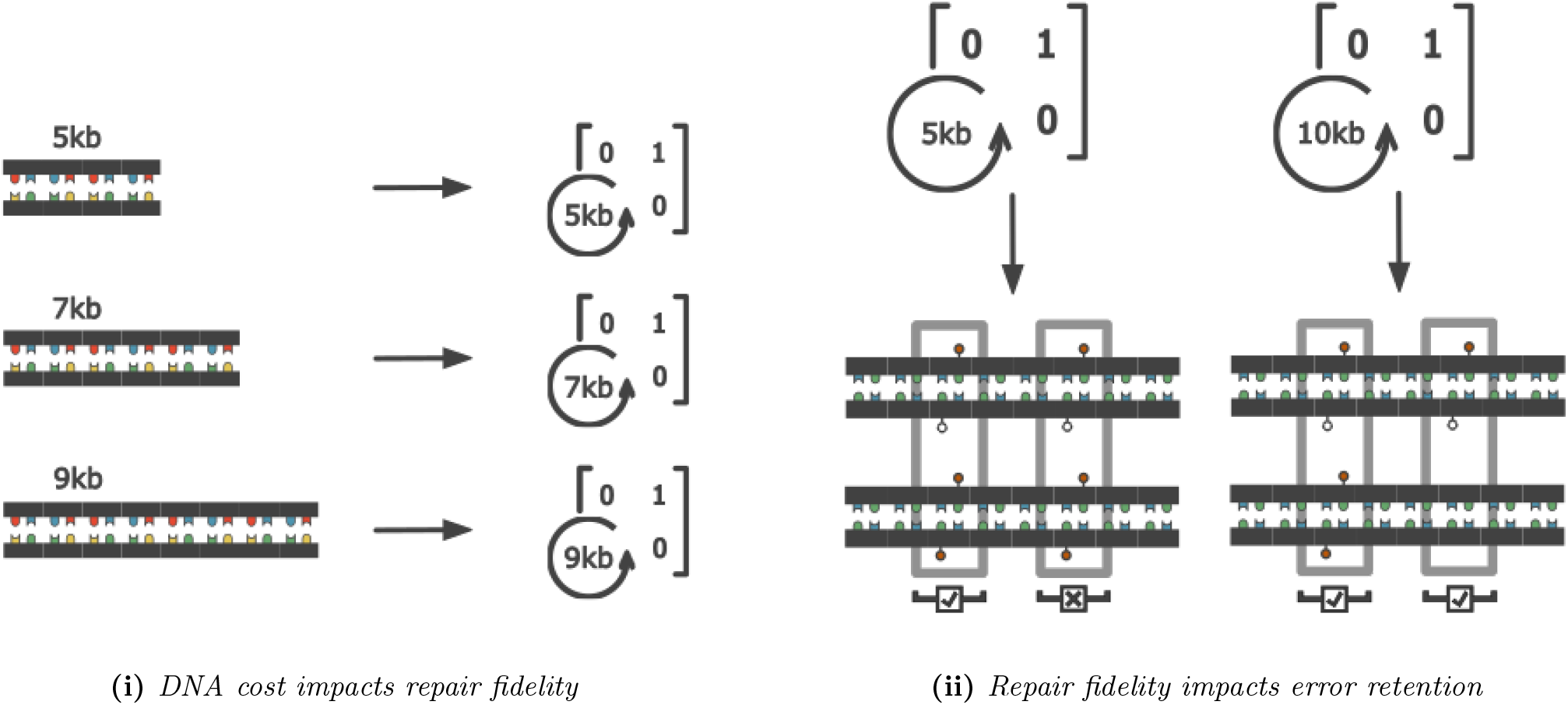
i. Any algorithm that corrects for epigenetic error must be comprised of proteins or RNA, which requires a region of DNA to encode. The size of the genes encoding the system will be related to the sophistication of the algorithm, but this will come at a survival cost commensurate to size. ii. The more sophisticated the algorithm for correcting error, the higher the fidelity to correct error, as more rules can be encoded to describe the correct state of a region based on more detectable environmental variables.

This is not to say that reconstruction is generally impossible, but that high-fidelity reconstruction is extremely informatically expensive and impossible to perform over a large number of cellular states.

As a result, an epigenetic algorithm reconstituting lost information would be forced to work within a spectrum between total lack of fidelity (randomly recreating data) and flawless fidelity, with the massive amount of information required for high fidelity restricting the majority of systems to error-prone reconstitution of damage.

### Legal Characters and Trust

When damage occurs in DNA it almost always produces a dictionary illegal character on one of the strands. In such cases (e.g., bulky adducts) DNA repair mechanisms can cheaply and effectively recognise that these new ‘characters’ in the DNA signal fall outside of the pre-defined set of legal dictionary characters: A, T, C, G. Both in this situation and when a dictionary legal character is created, DNA repair mechanisms must look for more information to determine which of the two strands to treat as an information backup of the original (Fig 1(i)). This is a system of trust and while imperfect, allows for the correct repair decision to be made the majority of the time. When the repair decision is incorrect, it might result in mutation: the introduction of an incorrect but dictionary legal signal element into the DNA signal. Damage can only be repaired in the context of a signal in which the damage is recognised as a dictionary illegal element. As mutation creates a legal character within the context of the signal representing the immediate DNA environment, there is not enough information on the original state of a dictionary legal signal to allow for entropy-neutral reconstitution of state, and so we can say that any dictionary legal error such as mutation is logically irreversible in the immediate context of repair enzymes. To repair this error in an entropy-neutral manner, the signal containing the error must be assessed in a higher syntax of which the local signal is but an element. We can say that the information scope must be broadened for repair.

DNA methylation sits outside the phosphate backbone and thus outside the system of trust which allows for limited local scope repair of DNA damage, and it has exactly two dictionary legal characters: fully methylated or unmethylated on both strands. Assuming no other information, this results in a situation where if one methylation is removed/added and a hemimethylated state is created, there is no logical way within the scope of a single repair enzyme to deduce which of the two legal characters the damage state originated from. The information of the original state is destroyed in the local scope, that which contains information limited to the methylation groups and immediately surrounding base pairs. We can therefore say that methylation damage is universally logically irreversible (as outlined in (18; 19; 20)) within the local scope of repair enzymes, with all the consequences of such a trait, namely obligate entropy increase upon repair (19; 21; 22). Put simply, all hemi-state DNA methylation created by damage is the equivalent to mismatched DNA bases with intact backbones and all epigenetic damage is consequently equivalent to mutation. We can say from this that epigenetic damage repair decision-making is a recognisable but not decidable language.

### Repair information is in the wrong place

In any situation of repair, the reconstitution of damage is limited by the amount of information available to the repairer. We can think of this as the scope of information that the repairer has access to. Any repair algorithm will sit within nested scopes of repair information: an enzyme might only be “aware” of the information in the immediate region of DNA it contacts; it has no access to information encoded in some distal section of DNA, or another cell, or another city. The super-entity of control represented by the system expressing and targeting that enzyme might well have access to a broader scope of information with which to target repairs. The caveat is that decisions about repair and consequently accurate repair can only occur within the scope of the information required and this may not be the scope in which the information exists. For example, the enzyme running along DNA has more up-to-date information about the current damage-state of the piece of DNA it sits upon than does the system that sent it to fix that damage. It does not always follow that the information scope of a subunit is a subset of that of a system with a broader scope. Information loss can result in logically irreversible damage within one scope and that same damage can be logically reversible within another. The question is: which scope has access to the information necessary to detect the information loss and which scope has access to the information necessary to repair the information loss? In non-mutational DNA damage, both of these sets of required information can exist within the same scope: that of the repair enzymes. In both DNA mutation and epigenetic mutation the information encoding the state of the ontological purpose of the governed system is exclusively found within systems that have access to information on ontological outcome. The information on the identity of the specific loci (base or CpG) under interrogation is limited to the repair enzyme while the repair enzyme remains at the locus of damage. This information is temporarily segregated from that of ontological outcome: the repair enzyme will have moved on and discarded information delineating location by the time the system is observably diminished in efficacy towards survival. It is therefore impossible to provide repair enzymes with the information necessary to correctly repair specific instances of mutation after the mutation has occurred, as well as any damage that could arise from more than one dictionary legal character. The information necessary for repair is not locally available at the point repair is locally possible.

### Ageing is the consequence of repair fidelity limitations

As there can be no mirroring backup to the epigenetic state and any algorithmic backup is limited below perfect fidelity, the logically irreversible information loss accrued within the epigenetic state will remain unrepairable in systems that have a complexity high enough that the information necessary for repair demands lossy compression. Damage will only be repaired up to the fidelity allowed for by the compression of repair. The only way to create logical reversibility in the systems and to reduce entropy is to increase the scope of the system until logical reversibility is possible.

When the amount of information necessary to create a system exceeds that which is beneficially storable in DNA, lossy compression will begin to be used as information is encoded in cellular context. This is the fidelity boundary: the point beyond which perfect fidelity is impossible. By storing information in the state of the local environment, systems can minimise the need to explicitly code functionality in DNA while retaining the information for approximate functionality, but with the consequence that they become logically irreversible as the low fidelity by which they are encoded results in multiple possible original states for the current system state. At this point, repair can occur but only in an entropic manner with a degree of error.

This fidelity boundary is never reached in simple systems but when systems expand in scope to allow for logical reversibility, they increase the information necessary for repair, approaching or crossing the fidelity boundary. If any system that influences the information necessary for its own repair is complex enough to demand definition past the fidelity boundary it will imperfectly repair itself when damage occurs, generating a feedback loop as it progressively repairs itself with decreasing fidelity. We suggest that simple systems are logically reversible below the fidelity boundary and complex systems influencing their own repair are not, inevitably becoming dysfunctional unless they are so valuable for organism survival that selection encodes the entire system within DNA. Only through the construction of a true logically reversible repair scope can the inevitably accruing system corruption be fully reversed.

When the information required for logical reversibility exceeds that storable in the immediate context of the cell, the scope must be extended again to allow for repair. Logical reversibility is then only achieved when the scope expands to include a known originator state, i.e. a stem cell. At this point, contextual algorithms with imperfect fidelity reconstitute the information of the cell (differentiation). In essence, the cell abandons its current state and returns to a point of known logical reversibility. Stem cells represent a type of cell that can be defined independently of context and thus in an informatically efficient manner. It is a simple, singular set of rules to encode, cheap and robust due to the lack of need to handle multiple definitions consequent to context. As epigenetic damage creates complexity not just in individual cells but in tissues and organs, the information defining the use of stem cells to reconstitute damage becoming itself progressively corrupted as tissue composition changes. This means that the scope that allows for logically reversible repair must be extended further back into epigenetic basality and more and more cells and eventually tissue discarded to allow for this. Discarding the information represented by these tissues and cells reduces the information necessary to encode a logically reversible state, and when this has occurred enough that identity can be defined below the fidelity boundary logical reversibility becomes possible. We suggest that this is the point that childbirth is the only solution available to the organism, in that the organism discards the entire body save for a single primordial stem cell that allows for the logical reversal of the entire organism. Through this lens, reproduction is not only a tool for selection but a necessity to shed age, and we argue this is why we, or the tissue that once was part of us, becomes young at the point of childbirth.

### Low fidelity creates error feedback

As epigenetic signalling fails, the systems governed by that signal will make incorrect decisions, resulting in a feedback cycle in which the epigenetic fidelity governing epigenetic fidelity fails, resulting in a recursive loss of epigenetic control as well as deregulation of all systems in which logical reversibility is impossible in the scope of repair. The deregulation of all cellular systems governed by epigenetic control is what, we suggest, gives rise to the phenotype of age and explains the generalised loss of cellular identity seen in ageing tissues.

Our theory suggests that ageing is itself the inevitable consequence of the impossibility of signal fidelity due to the specific dynamic of epigenetics being a single system in which it is impossible to design trust (through a mirrored backup) or an algorithm with perfect signal fidelity in systems where complexity is high enough that logical reversibility is impossible without crossing the fidelity boundary.

## 2 Results

### Methylation variance with age

The principles outlined above suggest that there is an inevitable accumulation of epigenetic damage with age, driving the structure of epigenetic signals into randomness. One measure of this dynamic is the progressing disparity between an individual’s DNA methylation loci with age. We obtained methylation data from externally generated datasets (outlined in Methods) and obtained from them beta values that represent the ratio of methylated to unmethylated measurements within each sample for each individual probe. The loci noise-to-age correlation (*nac*) for each dataset was obtained as described in methods. Results for datasets 1, 7 and 13 are summarised in Fig 3(i), 3(ii) and 4(i). All *nac* results and graphs for datasets 1-13 are provided in supplementary file 1. We used Benjamini-Hochberg correction to account for multiple testing, but most forms of multiple testing are heavily biased to extremely strong correlation, and in any analysis of stochastic noise the understanding of what represents ‘fluke’ correlation can be observed through the expectation that these will be represented by a normal distribution centered on 0 correlation.

**Figure 3:**
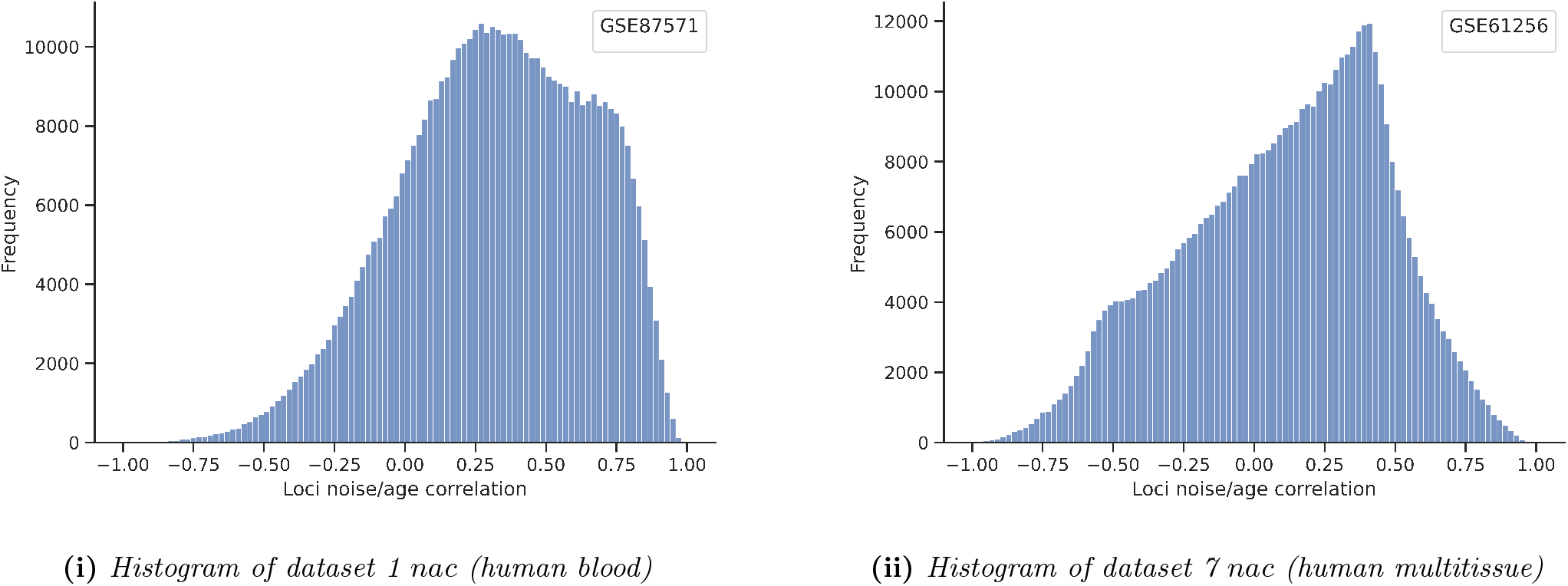
(i.) Dataset 1 (GSE87571 human whole blood) noise/age correlation (nac, as described in methods). Fluke correlation would be expected to be a normal distribution centered on zero. The peak at nac 0.75 represents a large population of CpG loci that increase in variance between samples with age. This dataset lacks a visible peak in the negative ranges of nac. (ii.) CpG SD correlation to age in Dataset 7 (GSE61256 human multitissue) nac. This dataset displays all three theorised populations, including a third peak within negative nac. We speculate that this peak represents genes switched on or off with age.

To decompose the *nac* of these tissues we looked to separate this central distribution from additional peaks representing subclasses of CpG with significant populations of strongly positive or negative *nac*. In all human datasets, save dataset 2 (where any such peak remains ambiguous and non visible due to the positioning of the fluke distribution), we see a unique peak of strong positive *nac*. In datasets 5, 6, 7 and 13 we observe a comparatively smaller peak of negative *nac* (dataset 6 being ambiguous due to the low number of samples within it above the age of 60). In non-human datasets the majority of loci approaching either -1 or 1 *nac*, likely consequent to there only being few recorded age groups in these datasets.

The general observation is that there are three approximate classes of loci: those that correlated negatively with age, those that correlate positively, and those representing ‘fluke’ correlation, centered on *nac* 0.00. For reasons discussed below, we term the population of CpG loci with abnormally high *nac* as ‘self-referential’ CpG/CGI (srCpG/srCGI), and the population of loci with a negative *nac* as ‘age-referential’ CpG/CGI (arCpG/arCGI).

We noticed that in almost all human datasets the central distribution, representing those CpG that do not belong to a class specifically relating to age, is slightly shifted most commonly towards positive correlation. The exception to this is dataset 13, comprised of 1400 samples (25 samples per year of age between 30 and 85) taken from a range of human datasets with multiple tissues being represented. This dataset was made consequent to the individual datasets analysis to see if independently of tissue type loci had a general trend towards high *nac*. The result (Fig 4(i)) was that the general peak representing typical loci became centred near a *nac* of 1, meaning that the typical loci accumulates noise in a manner that consistently correlates with age. This shift is likely consequent to the high number of samples and the decreased size of the age bins (from steps of 5 years to 1 year), allowing for more consistent estimation of *nac*. The better representation in this dataset of the upper age ranges is also likely a contributing factor. Due to the peak representing typical loci being centred near *nac* 1.0, the rightshifted peak seen in individual datasets is not visible. We believe this class of loci still exists, but is hidden by the increased *nac* of the general loci, and is more visible in the histograms of the individual datasets. It is also worth noting that in this higher fidelity dataset, the peak representing arCpG also remains intact, further arguing that these represent a unique class of CpG islands (CGI).

**Figure 4:**
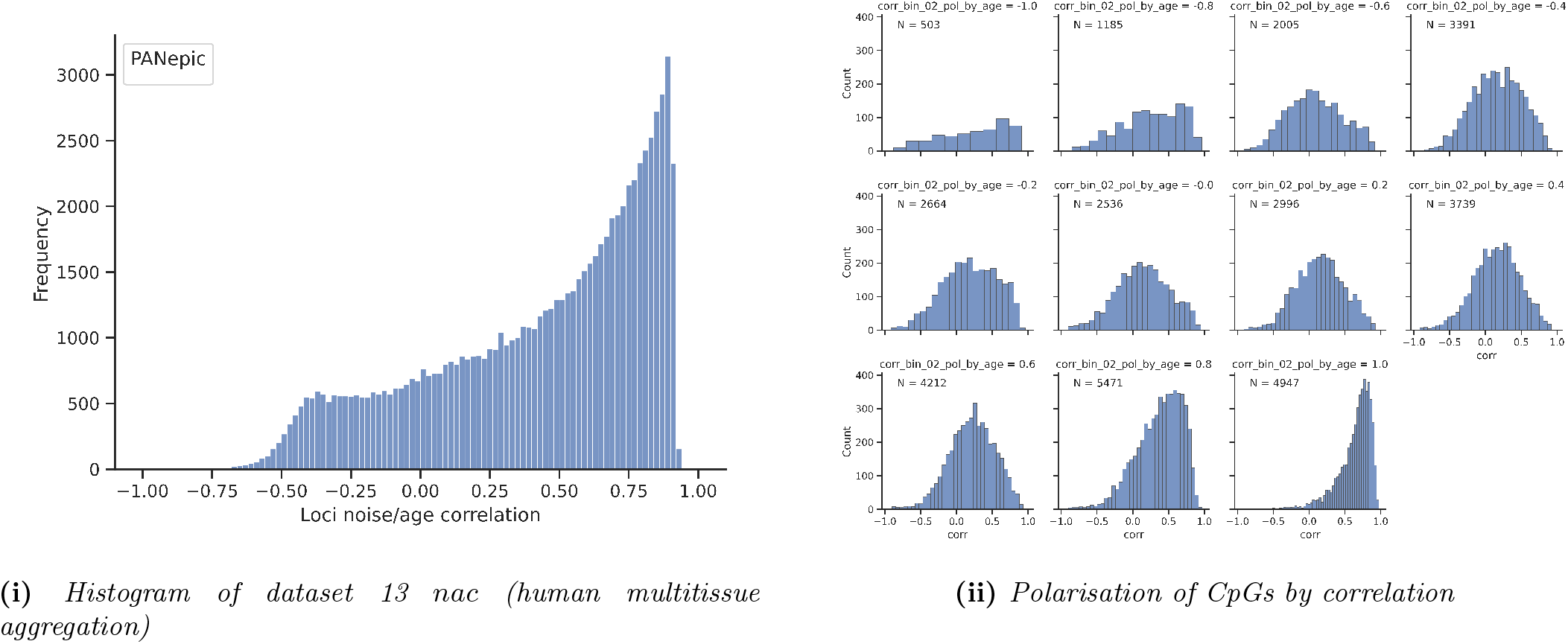
(i.) Dataset 13 (human multitissue aggregation) nac (as described in methods). Due to the larger number of samples, particularly in the higher age ranges where nac might be most expressed, and due to the increased fidelity of age measurement (values binned every 1 year as opposed to every 5 years), this dataset shows a clearer picture of typical CpG nac. With the exception of arCpG represented by the small peak around nac -0.4, it seems that all CpGs increasing in variance with age with a strength of correlation that results in the distribution of typical CpG nac beign centered on nac 1.00. We speculate that the smaller peak seen in the higher ranges of nac in individual tissue datasets is not visible on this graph due to the class being swamped by the majority position. (ii.) Dataset 1 (GSE87571 human blood) nac faceted by the correlation between beta value centralisation and age. CpG loci high nac are those in which beta centralisation increases with age.

### Methylation polarisation with age

We theorised that those CpG with high *nac* represented a class of noise-retaining loci (later termed srCpG). We expected the peak at r=0.00 to represent the *nac* of typical CpGs, and we next set out to characterise the nature of the negative correlations. We suggest that as an individual gets older there is a tendency to switch on or off genes as a mechanism to control noise. We expected that these genes would therefore tend towards polar beta values in their regulating methylation (either fully methylated or unmethylated). To explore this, we segregated *nac* by the polarisation of loci mean beta value with age in datasets 1 (Fig 4(ii)), 6, 10, and 14. This demonstrated that in these datasets the distribution of negative correlation is heavily skewed towards polarising loci, and that the peak representing srCpG becomes segregated to those CpG in which centralisation of beta value increases with age. This suggests that those loci with that decrease in variance with age indeed represent those genes that are increasingly regulated in response/consequent to age, and thus a different class of loci to those that, free of regulation, drift into variance.

### Methylation topology is an activation function

We hypothesised that if different methylation regions represented different classes of epigenetic control resulting from the need for discrimination in the amount of noise gene functionality was exposed to, evolution’s naturally conserving effect would unify these classes into a few different regulatory activation functions governed by a simple set of arguments. Were this the case, we would expect to observe conservation between the way specific loci were themselves regulated and therefore regulated the underlying gene. To visualise this, we performed Euclidean k-means clustering on sorted preparation of datasets 1 (Fig 5(i)), 6 10 and 14. In each of datasets 1, 6 and 10 (human and zebra, both longer lived mammals) we observed a definite grouping in locus topology, indicating that there are a few archetypical “activation functions” to which all DNA methylation belongs, and by which all methylation is regulated. It appears that DNA methylation is controlled by two types of functions, a linear function and a step function, which themselves act as components for a small range of combination activation functions. We classify these overall activation functions as linear functions, single-step functions, multistep functions, and “ragged” functions (functions containing regions with numerous fractional subpopulations independent of the majority ontological state). The topology of dataset 14 (GSE120137 mouse tissues) differs from that of longer-lived mammals (Fig 5(ii)). Both linear and step functions are observable, but the banding effect seen in human methylation data is absent.

**Figure 5:**
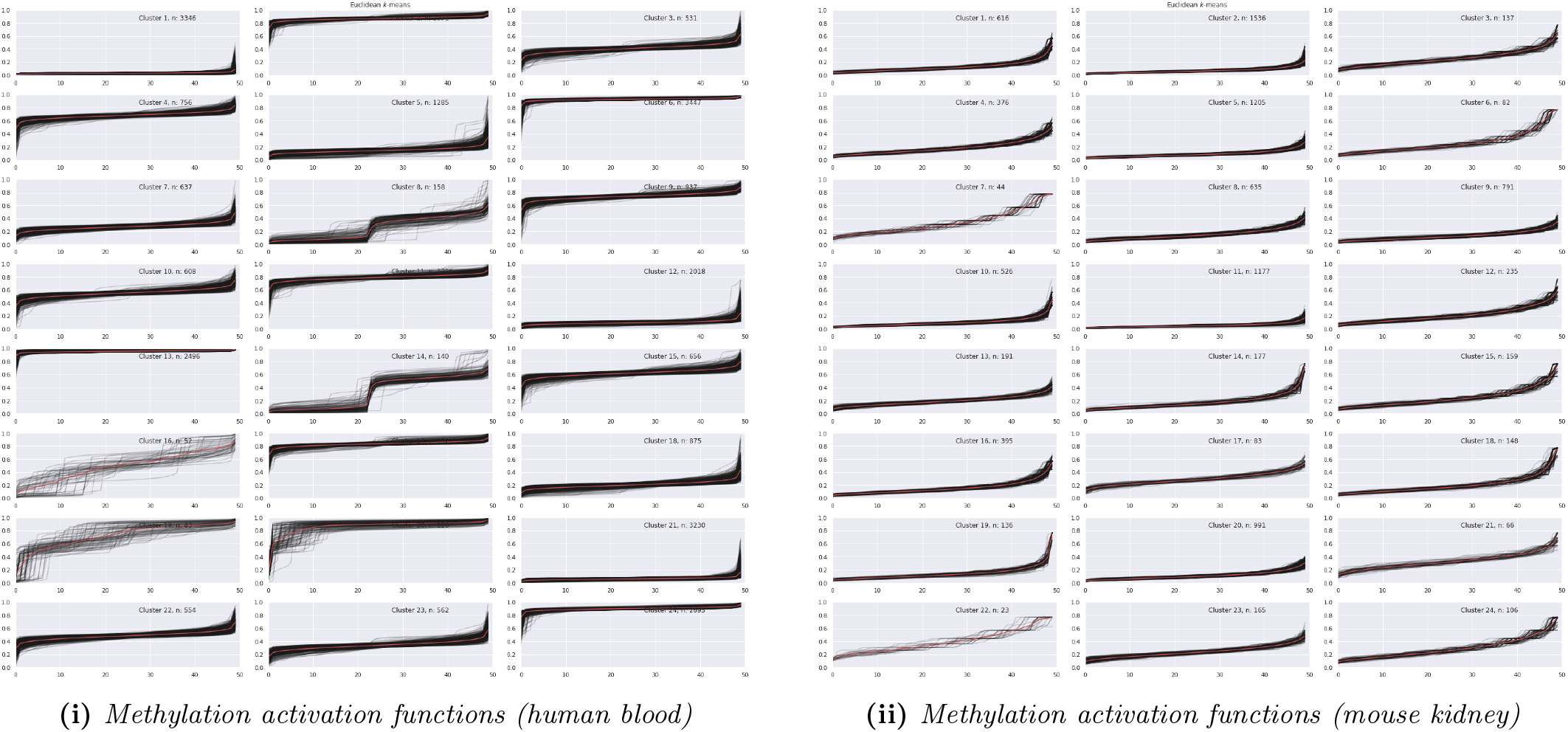
i. Topological activation functions of DNA methylation in dataset 1 (GSE87571 human whole blood). Sorted dataset clustered using Euclidean k-means. ii. Topological activation functions of DNA methylation in dataset 14 (GSE120137 mouse kidney). Dataset were sorted by value and clustered using Euclidean k-means (x resampled to 50).

Examining the topology of the clusters and individual CpG loci highlights that in most cases there is a majority position output by the function controlling methylation for each individual CpG locus. We assume that these are the positions for the ontological purpose of system function. In a large number of loci, there is also a ‘tail’ region of rapidly increasing or decreasing methylation approaching the nearest occupancy absolute (beta 0.00/1.00). These tails have the distinguishing feature of being an approximately consistent fraction of the total population size, but the difference between the tip of the tail and the mean beta value can vary substantially. We theorise that these tails represent a failure of control, in which a methylation area that is ideally at a given level of methylation loses its ability to regulate itself, resulting in mean betas that differ greatly from the ontological target. Mouse topologies increase at a comparatively steep and smooth linear rate with very little evidence of the “tailing effect” seen in the loci of longer-lived mammals (Fig 5(ii)), which suggests that there is comparatively less ‘failure of regulation’ because there is less regulation in the first place, mice not selecting as strongly for epigenetic fidelity as mammals more exposed to epigenetic mutation through lifespan and lack of predation. It is notable that these tails do not correlate to age in all loci. We believe that these may in some cases represent a loss of control in disease states and/or systems that lack the ability to create positive feedback into further epigenetic deregulation.

### Neural Networks can predict age from noise

We trained a Keras regressor on datasets 1 (Fig 6(i)), 4-6, 9-12, using VIP preprocessing on each, as outlined in methods. The purpose of this preprocessing was to eliminate any non-variance information in the dataset, giving the neural network no clue as to the overall beta values. All the neural network had access to was the knowledge of how relatively varied the beta values were in the age group the sample belonged to compared to the other age groups in the feature. Features were selected by random sampling and the results represent the mean of a Kfold split (f=5). Network definition and results are summarised in supplementary file 2. We observe that in all cases the VIP clock can predict age with great accuracy, illustrating that neural networks have the capability to predict based entirely on the amount of variance in a CpG site, rather than information on actual methylation levels.

**Figure 6:**
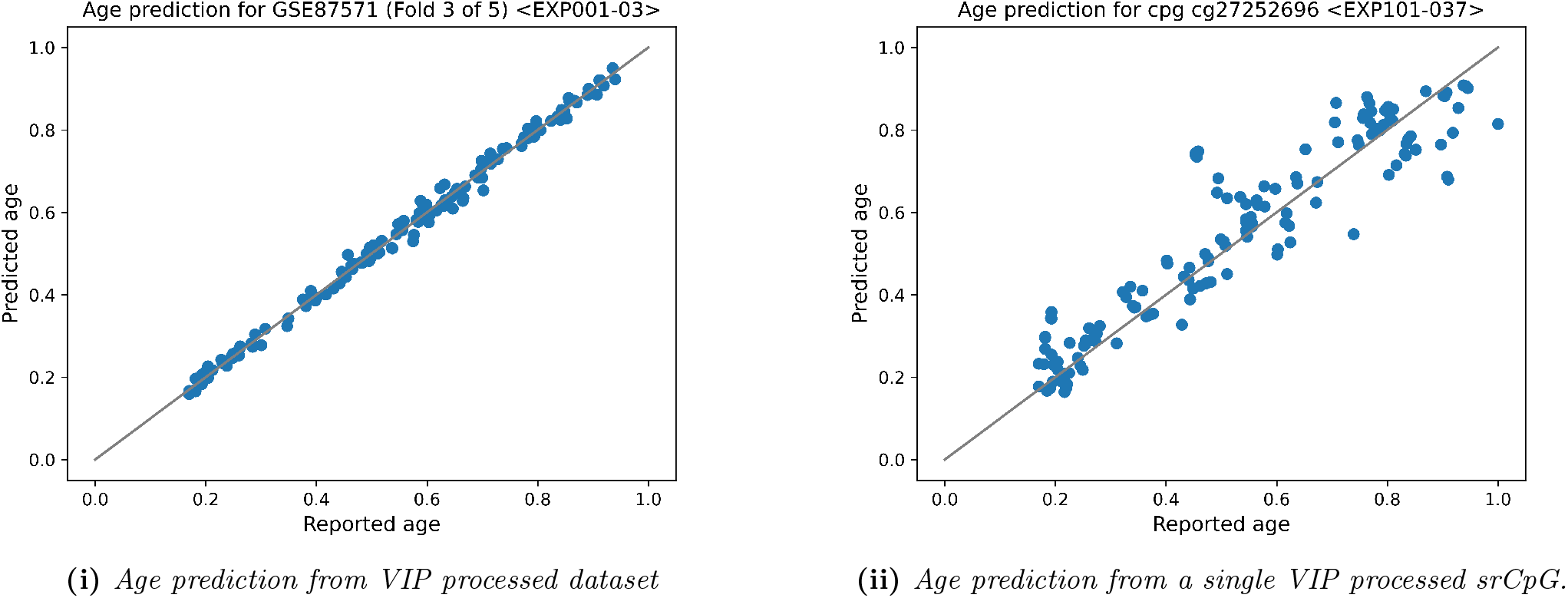
(i.) Regressor trained on 10000 DNA methylation loci beta values from dataset 1 (GSE87571 human blood). Chronological age (y) is measured against vs predicted age (x), with perfect prediction being represented by the samples in which the predicted age matches the chronological age. (ii.) An identical regressor trained on a single srCpG. N.B. Graphs correspond to results given in supplementary table via ‘EXP’ code.

To further demonstrate the capacity of such preprocessing, we selected twenty loci from dataset 1 with a *nac* of over 0.95, and used these samples with VIP preprocessing to construct single CpG networks (Fig 6(ii)). In all cases, these single CpG clocks were predictive of chronological age better than random, with a mean error of 9.18 years.

### Control is lost in the regulation of regulation

To improve the classification of high *nac* loci and to isolated proposed ‘srCGI’, we reasoned that ‘fluke’ correlation was more likely to be represented in individual CpG loci than entire CGI which should be more consistent in their ontological effect. To this end, we grouped loci by their CGI and calculated the SD of the correlations within these CGI.

We subset datasets 1, 2 and 13 to only include CpG from CGI with a *nac* SD below 0.15, and further subset to contain only CGI in which both mean CpG *nac* and centralisation (inverse of polarisation) were above 0.8. We then used methylGSA (23) to perform gene set analysis on unique genes associated with these loci and compared them against both those loci that fall outside this criteria and ten sets of randomly sampled CGI of equivalent size.

This preparation resulted in the enrichment of sequence-specific DNA binding (GO:0043565), mRNA Binding (GO:0003729), Ubiquitin Binding (GO:0043130), Methylated Histone Binding (GO:0035064), Chromatin DNA Binding (GO:0031490), RNA polymerase II transcription regulatory region sequence-specific DNA binding (GO:0000977), cis-regulatory region sequence-specific DNA binding (GO:0000987) and other terms that relate to transcriptional regulation (summarised in supplementary table 3), traits absent from randomly sampled control sets and the low polarisation set. It is interesting that those CGI with highest *nac* are those that are involved with the regulation of promoters and enhancers, i.e. the regulation of regulation.

## 3 Discussion

The role of epigenetic changes in ageing has been a major research focus, in particular since the discovery of epigenetic clocks. In this work we set out to break down the nature of epigenetic damage and characterise biological ageing as a failure of repair fidelity. To do this, we began by showing that chronological age generally correlates to the progressive dispersion of loci methylation state, and that there are two classes of loci (arCpG and srCpG) that significantly differ from the typical loci in how much the noise they accumulate correlates to age (Fig 4(i)). We can also demonstrate that these are distinct classes of CpG loci in how they are regulated in response to age (arCpG/arCGI) (Fig 4(ii)) and demonstrate through the ranked variance clock that age can be predicted exclusively from the knowledge of srCpG deregulation (Fig 6(i)), which we confirmed in a range of tissues and species. This fits exactly with our initial theory: epigenetic damage is inevitable due to the impossibility of mirrored backup and the bounding that limits any algorithmic repair to imperfect fidelity, and that damage causes and therefore correlates with age. We suggest that the distinct class of age-response CpG (arCpG) represent those genes that are being regulated increasingly in response to the phenotype of age provoking damage (senescence, DNA repair, stress response, etc). These loci are age referential because they react to age, switching genes on or off as part of the cellularstatic response to the deleterious effects of age. They have negative *nac* because they consequently become less noisy as biological age increases: a gene in a relatively unregulated state with range of potential outputs becomes tightly regulated, switched on or off and the regulatory elements becoming more tightly bound in value due to the pressures exerted. We suggest that srCpG represent those systems in which control fails in such a manner it has a knock on effect on other systems, the ground zero for failure, and argue that there is evidence that the reason for this subclass existing is that these srCpG control regulation itself, self-referential in their definition of state. Because srCpG cannot be provided with a system of trust and backup, as described in the theory above, they are doomed to fail, and take everything they control with them. As they eventually control all other regulation, they erode the general capacity of the cell to regulate epigenetics, evidenced by the typical shift of the general loci population towards positive *nac*.

We also find evidence for the progressive noise accumulation in the manner by which individuals deviate from the output of methylation activation functions. We suggest that there are ‘activation functions’ of regional epigenetic control that are observable through the limited range of outputs that define average methylation across populations (Fig 5(i)). These activation functions are defined through control of local methylation pressure, upon which will be applied mechanisms that we suspect to be a handful of evolutionarily conserved behaviours repeatedly applied in combination to produce a limited range of robust behaviours.

Within these activation functions, we suggest that deviation from the ontologically intended output of the activation function represents epigenetic damage, eventually resulting in deregulation of the governed gene expression. It seems likely that this is what is represented by the topology ‘tails’, samples in which loci control has either become aberrant (or possibly some cases where the loci were measured during a transition state).

We propose that this epigenetic damage would result in a feedback cycle, in which deregulation would lead to further deregulation through the disruption of maintenance and repair of the epigenetic regions, and to the phenotype of age through the general deregulation of cellular systems.

This would fit the profile of ageing as a robust, gradual process, with slow, reliable progress made as deregulation accumulates, accelerating toward network failure as the feedback cycle picks up pace. We can see in the gene ontology results (supplementary table 3) that in all organisms and tissues, those genes regulated by the loci in which deregulation correlates to age are genes governing promotors and enhancers.

We suggest that this is because promotors and enhancers have a unique feature that precludes polarising their regional control for regulation: they need to regularly reconfigure the local methylation state consequent to the current state of transcription. We suggest this makes them self-referentially defined, in that the definition of the epigenetic signal of a promotor/enhancer modulator relies in part on its own current state (such that any damage results in damage to any rule from which the signal could be corrected), and thus representing a class of loci in which epigenetic regional control cannot be correctly defined once epigenetic damage has occurred. We suggest damage accrues in these regions and the global deregulation of transcription that occurs consequent to this gives rise to the general phenotype of age.

Our work characterises the cell as incapable of perfect reconstitution of all instances of epigenetic damage, and that it must instead pick and choose which systems to hold to high fidelity and which to allow low fidelity. In some cases, we argue, low fidelity is forced either way. There can be no perfect fidelity backup of all epigenetic information stored in the cell.

We would argue against the idea that any specific cellular subsystem explains the general dynamic of age, rather suggesting that the frequency of epigenetic events is likely to be the main influence of the rate of epigenetic damage and it is a core dynamic of epigenetic control that information loss must occur in the regulation of any adaptive system. The nature of the systems themselves is irrelevant to the frequency of epigenetic modification they demand and the fact that such modification must in these systems be self-referentially defined. We argue that DNA repair and other correlated behaviours are simply representatives of this class of epigenetic regulation and those models that recognise the link between age and particular systems such as DNA repair (12; 11) are measuring the rate at which epigenetic modification within these systems leads to epigenetic damage. It has been demonstrated that there is a general stochastic loss of methylation over time (24) and that hemimethylated states can spontaneously occur through methyl group drop-off or enzymatic error (25), and we suggest that this might provide a base rate at which biological ageing progresses. We suggest that the idea the cell has a problem with information loss is too broad; information is lost all the time. We suggest that the real problem is the limit of fidelity that biological compression must labour under, the necessity for the use of lossy compression and a top-level system of information handling that simply cannot be designed in a manner with sufficient damage recovery. Information is lost in systems that cannot correct for information loss, and the only type of system that fits that criteria are systems that have no external oversight to provide logical reversibility and must therefore maintain their own identity. The issue is not the general occurrence of information loss, but the existence of information loss in specific systems, those incapable of identity reconstitution through the necessity of self reference. We argue noise in the identity of noise identification systems is the root cause of ageing.

This theory, therefore, provides a full line of reasoning from the logical necessity of epigenetic damage through to the phenotype of age. The outlined theory is not limited to methylation-based damage: the duplication of histone markers and damage within that system shares the exact same issue as methylation, as will any outer layer system of information fidelity. In an effort to reduce the information necessary to maintain a system in which damage is logically reversible, we propose mammals use childbirth to shed complexity, reducing the information necessary to a single cell. In primordial cells, stem cells and the simple cellular systems of organisms without epigenetic regulation, damage is reversible through the encoding of state in algorithms that can define the correct informatic state within the genetic code. This is possible due to the ability to prescribe what a cell should be independently of its environment, and results in logical reversibility. We suggest that logical reversibility is a necessary objective for any organism existing in a noisy channel.

## 4 Methods

### Datasets

Datasets were handled in Python (v3.10.6) and processed using Pandas (v1.5.3), Numpy (v1.22.4), methylprep (v1.7.1) and GEOparse (v2.0.3). Datasets were visualised with a combination of Matplotlib (v3.1.3) and Seaborn (v0.11.2). Datasets were processed from IDAT using methylprep default function (steps = [“all”]) or obtained by direct download from the NCBI GEO repository (https://www.ncbi.nlm.nih.gov/). Datasets were acquired with the most recent version as of 01/09/2022. In cases of a dataset containing multiple tissues, the tissue with the greatest number of samples was used unless specified in supplementary file 1. The datasets below, tissues selected and the processes done to them are described in supplementary file 1. In all cases, columns with na values were removed prior to further processing.

- Dataset 1: GSE87571 (human blood) (26)
- Dataset 2: GSE132203 (human blood) (27)
- Dataset 3: GSE116339 (human blood) (28)
- Dataset 4: GSE183920 (human white blood cell) (29)
- Dataset 5: GSE175458 (human lung) (30)
- Dataset 6: GSE41826 (human brain) (31)
- Dataset 7: GSE61256 (human liver, muscle and adipose) (32)
- Dataset 8: GSE183647 (human brain tumor) (33; 34)
- Dataset 9: GSE208713 (human liver tumor) (35)
- Dataset 10: GSE184223 (zebra blood) (36)
- Dataset 11: GSE184216 (roe deer blood) (37)
- Dataset 12: GSE164127 (bat skin) (38)
- Dataset 13: Aggregation of datasets 2-5, 7-9
- Dataset 14: GSE120137 (mouse multi-tissue) (39)

### Dataset Processing

These datasets were used as a base to create additional datasets using the following transformations, referred to by the code in brackets:

### SD by age (SbA)

Samples were binned into groups covering five years of age (e.g. 60-65 years old). Within each bin, the standard deviation was taken for the beta values within each individual CpG loci, resulting in a table of age group SD by loci. Bin population was restricted to the lowest common bin population above 8, resulting in bins of equivalent size.

### Variance Isolating Preprocessing (VIP)

Variance Isolating Preprocessing (VIP) datasets were created by sorting samples by age and running a five-row sliding window on each CpG taking the SD for each window step. The age of each step was the mean age of the samples used for the window. The dataset then had all values replaced by a rank indicating their position from lowest number to highest. This was done to remove all possibility that the loci SD provided to the network was allowing it to estimate beta values. Samples were then shuffled.

### Polarisation by age (PbA)

The polarisation datasets were created by modifying the initial dataset with the following transformation on every value (*x*):

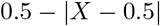

Samples were binned into groups covering five years of age (e.g. 60-65 years old), obtaining the mean and dropping any bin that did not have at least n=5. These values represented the final dataset.

### Statistical Tests

Statistical tests were performed using scikit-learn (v1.0.2) and Benjamini-Hochberg multiple correction was handled using the ‘multipletests’ function from statsmodels (v0.12.2). Beta plots were resampled using ‘TimeSeriesResampler’ and clustering was performed using ‘TimeSeriesKMeans’ from the package tslearn (v0.5.3.2). A random seed of 0 was used for all clusterings. Pearson’s R correlation was performed with the pandas ‘corrwith’ function, feature ranking and sliding windows were performed using the pandas ‘rank’ and ‘rolling’ functions respectively, and feature sorting was performed with the Python built-in ‘sorted’ function.

### Noise to Age Correlation (*nac*)

For each CpG, we binned samples into age groups as described in supplementary table 1 and took the standard deviation for each bin. We then performed a Pearson’s R correlation between the age-binned SD and age resulting in the noise-to-age correlation (*nac*) for the CpG.

### Deep Learning and State Machine

Deep learning was performed using Tensorflow (v.2.11.0) Keras. The methylation state machine was coded in Python (v3.8.10).

### Deep Learning preprocessing

Processing obeyed the following order. Sample and feature order were randomised. Any validation set was isolated. Special preprocessing, such as VIP was performed. Any post-preprocessing validation set was isolated. The network was then run at k-fold (5).

### Gene Ontology

GO analysis was performed using the Enrichr tool (40). Datasets used were subset to those CpG with known CGI identities and then individual CGI values were obtained for mean *nac, nac* SD and mean PbA, which were then used for subsetting the datasets. For each dataset, the background used was all the possible genes associated to the dataset when subset for known CGI identity (all genes that could have been selected had the CGI dataset been included in it’s entirety.) For each dataset, 10 random sample populations of equivalent size to the hypothesis subset were generated alongside the hypothesis subset (those CGI where mean *nac* was over 0.80, *nac* SD was below 0.15 and PbA was below 0.20), as well as a population comprising all of the CGI that fall outside the hypothesis set.

## Data Availability

- Supplementary file 1: Dataset descriptions and *nac* result graphs
- Supplementary file 2: Clock results
- Supplementary file 3: GO enrichment results

## With thanks to…

Ali Doğa Yücel provided support and discussion for the presentation of the theory and copyediting of the paper.

Paul Ka Po To provided optimisation for the neural network and discussion of normalisation techniques.

Work in our lab is supported by grants from the Wellcome Trust (208375/Z/17/Z), Longevity Impetus Grants, LongeCity and the Biotechnology and Biological Sciences Research Council (BB/R014949/1 and BB/V010123/1).

Conflicts of interest: JPM is CSO of YouthBio Therapeutics, an advisor/consultant for the Longevity Vision Fund and NOVOS, and the founder of Magellan Science Ltd, a company providing consulting services in longevity science.

