## Supplementary table 2 for "Epigenetic fidelity in complex biological systems and implications for ageing"

**Supplementary table 2.1 (network structure):**

Optimizer = Adam, learning rate = 0.001

Normalizer
Dense (256), elu activation, 0.001 regularisation
Dropout (0.4)
Dense (256), elu activation, 0.001 regularisation
Dropout (0.4)
Dense (128), elu activation, 0.001 regularisation
Dropout (0.4)
Dense (128), elu activation, 0.001 regularisation
Dropout (0.4)
Dense (1)

**Supplementary table 2.2 (network results):**

GSE87571 human blood sliding window dataset

| loss | mean_absolute_error | mean_absolute_percentage_error | mean_squared_error | mean_squared_logarithmic_error |
|---|---|---|---|---|
| 3.278847 | 2.49334 | 5.781902 | 14.78427 | 0.007394 |
| 2.910811 | 2.146641 | 4.316648 | 9.887054 | 0.002963 |
| 2.789407 | 2.030376 | 4.284602 | 7.567978 | 0.002919 |
| 3.96432 | 3.213025 | 9.093208 | 25.352938 | 0.022354 |
| 3.265279 | 2.485748 | 4.974205 | 12.412249 | 0.004005 |
| 3.054671 | 2.323184 | 6.811276 | 22.292887 | 0.018142 |
| 3.49524 | 2.72506 | 5.633029 | 13.04577 | 0.004682 |
| 3.123314 | 2.383826 | 5.058978 | 13.843047 | 0.006119 |
| 2.879027 | 2.099964 | 4.697271 | 7.068963 | 0.003028 |
| 2.801393 | 2.052816 | 5.083001 | 7.088758 | 0.004939 |

GSE41826 human brain sliding window dataset

| loss | mean_absolute_error | mean_absolute_percentage_error | mean_squared_error | mean_squared_logarithmic_error |
|---|---|---|---|---|
| 6.499018 | 5.538018 | 13.792947 | 56.430656 | 0.026553 |
| 4.526168 | 3.565236 | 9.850774 | 22.084307 | 0.013651 |
| 4.742898 | 3.774138 | 11.075229 | 26.10886 | 0.018946 |
| 4.930711 | 3.980876 | 11.641271 | 27.958364 | 0.020088 |
| 4.528511 | 3.575025 | 10.189425 | 19.763241 | 0.014108 |
| 4.905636 | 3.943056 | 11.787901 | 22.967678 | 0.018282 |
| 4.244316 | 3.281458 | 8.75231 | 19.262445 | 0.01164 |
| 7.277723 | 6.318108 | 16.905029 | 77.715202 | 0.044579 |
| 5.968963 | 5.008518 | 13.219999 | 48.412685 | 0.025884 |
| 6.409532 | 5.46239 | 11.937749 | 52.96394 | 0.01888 |

GSE87571 human blood ranked dataset

| loss | mean_absolute_error | mean_absolute_percentage_error | mean_squared_error | mean_squared_logarithmic_error |
|---|---|---|---|---|
| 2.806146 | 1.990457 | 4.464902 | 8.621114 | 0.003668 |
| 3.777213 | 2.985807 | 6.346775 | 19.10166 | 0.008602 |
| 3.022134 | 2.228419 | 5.82268 | 7.766428 | 0.005536 |
| 3.797386 | 3.012566 | 8.278977 | 31.108824 | 0.022955 |
| 4.463407 | 3.692132 | 7.345172 | 34.811459 | 0.010868 |
| 3.233589 | 2.428051 | 5.272555 | 11.317239 | 0.00451 |
| 3.7384 | 2.932013 | 6.420062 | 16.755358 | 0.009854 |
| 3.985407 | 3.194638 | 7.950692 | 20.701326 | 0.014227 |
| 3.456521 | 2.651514 | 5.858628 | 13.930508 | 0.006335 |
| 3.421491 | 2.635275 | 6.420143 | 12.805676 | 0.006371 |

GSE41826 human brain ranked dataset

| loss | mean_absolute_error | mean_absolute_percentage_error | mean_squared_error | mean_squared_logarithmic_error |
|---|---|---|---|---|
| 7.058796 | 6.092673 | 14.992749 | 58.813419 | 0.032072 |
| 5.162893 | 4.194484 | 11.599635 | 29.919556 | 0.020083 |
| 4.402566 | 3.438444 | 10.672811 | 16.382401 | 0.015853 |
| 5.440671 | 4.477244 | 13.087822 | 26.685974 | 0.021866 |
| 4.407511 | 3.451558 | 9.466867 | 18.307917 | 0.013008 |
| 4.019556 | 3.062629 | 9.440925 | 13.215676 | 0.010728 |
| 6.586806 | 5.624051 | 14.127192 | 47.741512 | 0.025925 |
| 5.140876 | 4.190904 | 10.513003 | 28.663975 | 0.013623 |
| 6.985198 | 6.026913 | 14.87104 | 83.877747 | 0.034321 |
| 6.700014 | 5.747361 | 13.423739 | 73.267311 | 0.027034 |
